# Modeling Sympathetic Neuro-Cardiac Interactions in a hiPSC-Based Microphysiological System

**DOI:** 10.64898/2026.05.06.723218

**Authors:** Jean-Baptiste Reisqs, Yvonne Sleiman, Mohamed Boutjdir

## Abstract

The cardiac autonomic nervous system is a key driver of various cardiac disorders and arrhythmias. However, investigating neuronal regulation of the human heart has proven difficult due to immitted and reliable experimental models.

Here, we present a novel microphysiological system utilizing a compartmentalized microfluidic device (MFD) to integrate co-cultured human induced pluripotent stem cell (hiPSC)-derived cardiomyocytes (hiPSC-CMs) and sympathetic neurons (hiPSC-SNs). MFD is composed of two wide-open chambers separated by microfluidic microchannels.

hiPSC-SNs were characterized by confocal imaging and RT-qPCR for the expression of peripherin, tyrosine hydroxylase, and β-tubulin III, as well as high levels of dopamine β-hydroxylase and nicotinic acetylcholine receptors. Furthermore, patch-clamp techniques confirmed their functional maturity, showing spontaneous action potentials and positive responses to nicotine (1µM). Co-culturing hiPSC-CMs and hiPSC-SNs within the MFD facilitated axonal projection into the cardiomyocyte chamber, establishing a physical connection between the two cell types. After 10 days of co-culture, functional integration was confirmed by a significant increase in the action potential frequency and beating rate of hiPSC-CMs, as recorded by patch-clamp and video motion tracking, respectively. Notably, nicotine application in the neuronal chamber accelerated these rates in hiPSC-CMs chamber, whereas the administration of the β-blocker, propranolol (5µM), effectively decreased the beating rates. Collectively, these data demonstrate the feasibility of differentiating hiPSCs into functional sympathetic neurons and establishing a robust neuro-cardiac interface. This microphysiological system represents a powerful platform for investigating disorders characterized by impaired neuro-cardiac interactions, offering a valuable tool for both disease modeling and pharmacological screening.

## Introduction

The autonomic nervous system (ANS) includes of the sympathetic nervous systems, which innervates the heart^1,2^. The brain-heart axis is important because the ANS regulates heart rate, contractile strength and kinetics, and action potentials (APs)^3^. Specifically, the sympathetic nervous system (SNS) is responsible for the “fight or flight” response^4^. The SNS innervate the myocardium via a network of postganglionic nerve fibers which release norepinephrine via calcium-dependent exocytosis^5^. Sympathetic cardiac regulation involves a cholinergic to adrenergic relay^6^. Preganglionic fibers release acetylcholine to activate postganglionic nicotinic acetylcholine receptors. Once activated, postganglionic neurons secrete norepinephrine, stimulating cardiomyocytes β-adrenergic receptors to increase heart rate and contractility^7,8^. Under pathological conditions the increase of β-adrenergic signaling could trigger life-threatening cardiac events such as in inherited arrhythmia syndromes like the long QT syndrome or catecholaminergic polymorphic ventricular tachycardia (CPVT)^9–11^. Pharmacological interventions, based on β-blocker, or cardiac sympathetic denervation are preventive strategies for these types of arrhythmias, but the cellular basis underlying triggering of arrhythmias remain not fully understood, hence the need for co-culture induced pluripotent stem cells (hiPSCs) into sympathetic neurons (iPSCs-SN) and cardiomyocytes (iPSCs-CMs) organ-on chip microfluidic device (MFD) to characterize the brain-heart axis.

While in vivo animal studies have underscored the critical role of the ANS in cardiac homeostasis and trophic development, they face significant translational hurdles^12^. For example, the density and role if innervation in basal cardiac function, and general cardiac electrophysiology differ from humans and large animals^13,14^. Although large animal models offer closer physiological proximity, their utility for high-throughput mechanistic investigations is severely limited by prohibitive costs and logistic^15^.

Collectively, to address this translational gap, human induced pluripotent stem cell (hiPSC) technology has emerged as a powerful tool for the neuro-cardiac connection studies. The possibility to derivate these cells into hiPSC-CMs, and more recently into hiPSC-SNs from the same subject have opened new avenues of investigations and precision medicine^16–18^. The classic co-culture systems, where hiPSC-SNs and hiPSC-CMs are mixed in a single dish, present also their own set of limitations. Recently, an innovative compartmentalized systems with removable inserts, facilitate the strategic co-culture of hiPSC-SNs and hiPSC-CMs^19^. This approach enables initial isolation of each cell type, allowing an independent maturation, followed by insert removal to promote the development of functional synaptic connections^19^. However, like previous studies, the co-culture needs a mixing co-culture media, which compromises cell viability and/or leads to uncontrolled cellular phenotypes, introducing experimental bias^18–20^. In addition, the conventional co-culture in the same dish lacks the spatial organization required to distinguish between somatic and axonal signaling. Finally, pharmacology intervention becomes inherently difficult and cell-specific responses impossible to distinguish. Consequently, there is a need for a platform that maintains the human-specific advantage of hiPSCs, while providing a structural compartmentalization necessary to study the neuro-cardiac junction.

In this study, we develop a microphysiological system (MPS), based on the use of microfluidic device, to model the neuro-cardiac connection using hiPSCs. This MPS is composed by 2 wide open chambers connected through microchannels, which successfully create a functional connection between hiPSC-SNs and hiPSC-CMs.

## Methods

### hiPSC culture

Biological samples were handled in accordance with the Declaration of Helsinki with informed consent and standardized approved protocols by a local ethics committee (S14-00862). The protocol was approved by the Institutional Review Board Committee at the VA New York Harbor Healthcare System (1750139). hiPSCs lines were obtained from Stanford University Cardiovascular Biobank, from a female patient aged at 20 years old.

### Neuronal induction, differentiation of sympathetic neurons from hiPSCs

The hiPSCs were plated and cultured on hESC-qualified Matrigel^TM^ (Sigma-Aldrich, MO, USA) at 30,000-50,000 cells/cm^2^ with mTeSR^TM^ (StemCells Technologies, CANADA). For the sympathetic neuronal differentiation, we used a protocol previously published by Winbo and collaborators^18^. Briefly, the neuronal induction was started at 70% confluency of hiPSCs, and a series of small molecules were used to initiate the trunk neural crest cells (tNCCs) to sympathetic neurons precursors, and finally immature sympathetic neurons (protocol of neuronal differentiation is illustrated in **Figure 1**). The media was progressively replaced, by 25% increment from mTeSR to Neurobasal medium with N2 Supplement (Gibco, NY, USA). At the end of the neuronal differentiation process, the immature sympathetic neurons are dissociated and replated on Geltrex and Poly-D-Lysine-coated 35-mm dish or into the microfluidic devices in neuronal medium (Neurobasal Medium + N2 Supplement (Gibco, NY, USA) + B27 Supplement (Gibco, NY, USA) + 2mM L-Glutamine, 0.2mM ascorbic acid, 0.2cAMP, 10ng/mL NGF, 10ng/mL BDNF, and 10ng/mL GDNF) and BMP4. After 2 days, the hiPSC-SNs were cultured in neuronal medium, without BPM4, for 30 days for further neuronal maturation.

**Figure 1:**
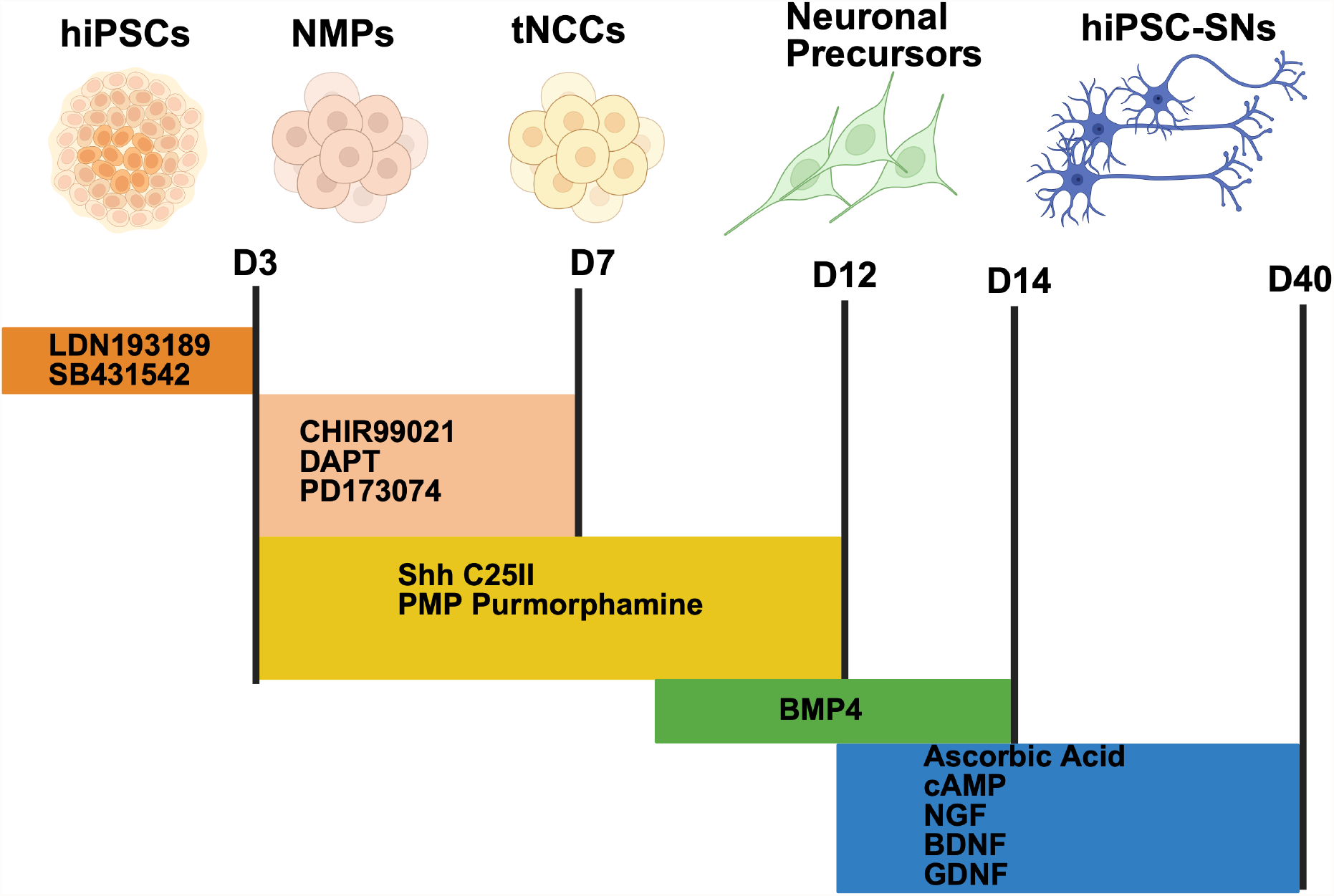
Schematic of differentiation protocol for sympathetic neurons-hiPSCs. hiPSCs at 70% of confluency were first treated with 500 nM inhibitor LDN 193189 and 10 µM SB431542 until day 3 (D3). Between D3 and D7, hiPSCs were then treated with 3 µM CHIR99021, 10 µM DAPT and 0.2 µM PD173074. In the meantime, between D3 until D12, hiPSCs were also treated with 60 ng/mL Shh C25II and 1µM PMP Purmorphamine. At D10, 10ng/mL of BMP4 were added. At D12, neuronal precursors were treated with 0.2 mM ascorbic acid, 0.2 nM cAMP, 10 ng/mL NGF, 10 ng/mL GDNF, and 10 ng/mL BDNF for further maturation. The hiPSCs media (mTeSR) were subsequently replace by Neurobasal medium (25% increments) at D4, D6 and D8. hiPSCs: human induced pluripotent stem cells; NMPs: neuromesodermal progenitors; tNCCs: trunk neural crest cells; hiPSC-SNs: hiPSC-sympathetic neurons

### Differentiation of cardiomyocytes from hiPSCs

In parallel of the sympathetic neuronal differentiation, the hiPSCs were differentiated into cardiomyocytes as previously described^21^. Briefly, the cardiomyocyte differentiation was initiated, at 90% of confluency, with STEMdiff^TM^ Ventricular Cardiomyocyte Differentiation Kit (StemCells Technologies, ON, CANADA). At day 8 after the initiation, spontaneous beating could be observed and the hiPSC-CMs were cultured with STEMdiff^TM^ Cardiomyocyte Maintenance medium (StemCells Technologies, ON, CANADA) for 30 days for further maturation.

### hiPSC-SNs and hiPSC-CMs co-culture

For the hiPSC-SNs and hiPSC-CMs co-culture, we used MFD OMEGA device (eNUVIO, Québec, CANADA). This device is composed by 2 open chambers connected by microfluidic microchannels (**Figure 4**). The unique open chamber design offers the advantage to perform patch-clamp and video-recording experiments. This microfluidic device uses asymmetric media volume (150 µL in neuronal compartment and 100 µL in cardiomyocytes compartment) to promote axonal outgrowth through the microchannels into the adjacent compartment, while maintaining fluidic isolation of the two chambers. The day before hiPSC-SNs seeding, MFD was coated with a solution of Poly-D-lysine (100µg/mL, Gibco), incubated overnight at 4°C, and rinsed 3 times with sterile water. Then, 3 hours before seedings, MFD were coated with Geltrex 1X (Gibco). Newly differentiated hiPSC-SNs were then seeded into the neuronal compartment in neuronal medium supplemented with BMP4 (10ng/mL) for 2 days. The hiPSC-SNs were maintained for 30 days for further maturation and start to form neurites and projecting axon into the microchannels and invaded the cardiomyocyte compartment. After this period, hiPSC-CMs aged 30 days were seeded into the cardiomyocytes compartment of the MFD. The ratio used for the co-culture was 10:1 (200,000 hiPSC-CMs to 20,000 hiPSC-SNs per MFD). After 10 days of co-culture, the experiments were performed.

### Immunohistochemistry and Imaging

For immunofluorescence, hiPSC-SNs and hiPSC-CMs are first fixed with 4% paraformaldehyde and permeabilized in a PBS solution containing 0.1% Triton X-100 for 15 min. The cells were saturated using Normal Goat Blocking Buffer (Elabscience, TX, USA) for 30 min. The cells were stained for 24 hours at 4°C using PBS containing primary antibodies. The secondary antibodies, goat anti-mouse IgG Elab Fluor 488 (1:200, Cat# E-AB-1056, Elabscience, TX, USA) and goat anti-rabbit IgG Elab Fluor 594 (1:200, Cat# E-AB-1060, Elabscience, TX, USA) were then incubated for 2 hours at 4°C in the dark. The nucleus was labelled using DAPI Reagent (Cat# E-IR-R103, Elabscience, TX, USA) for 5 min. Cells were observed at 63x objective using a Zeiss LSM800 confocal microscope (Zeiss, NY, USA) and examined with ZEN software.

The primary antibodies used in this paper were: mouse anti-alpha actinin (1:200, Ca# A7732, Sigma, MO, USA); mouse anti-Tyrosine Hydroxylase (1:200, Ca# AB152, Sigma, MO, USA); rabbit anti-beta-3 Tubulin (1:200, Ca# 86069, Invitrogen, CT, USA); rabbit anti-Peripherin (1:200, Ca# AB1530, Sigma, MO, USA).

### Quantitative reverse transcription polymerase chain reaction

RNA extraction from hiPSC-CMs and hiPSC-SNs were isolated using a NucleoSpin RNA kit (Macherey-Nagel, PA, USA) followed by a reverse transcription to obtain cDNA using the Transcriptor Universal cDNA Master Mix (Sigma-Aldrich, MO, USA), according to manufacturer’s’ protocols. For detection of the gene of interest, cDNA was amplified by the PowerUp SYBR Green Master Mix (Applied Biosystems, CA, USA) for RT-qPCR. cDNA amplifications were performed with the Quant Studio 5 from Applied Biosystems. Human Large Ribosomal Protein (RPLPo) was used as a reference gene. The genes of interest studied by RT-qPCR included *Tyrosine hydroxylase (TH), Dopamine* β*-hydroxylase (DBH)*, β*-3 Tubulin (TUBB3)*, and *α4 subunit of the neuronal nicotinic acetylcholine receptor (CHRNA4)*. List of Primers are found in **Table 1**.

**Table 1:**
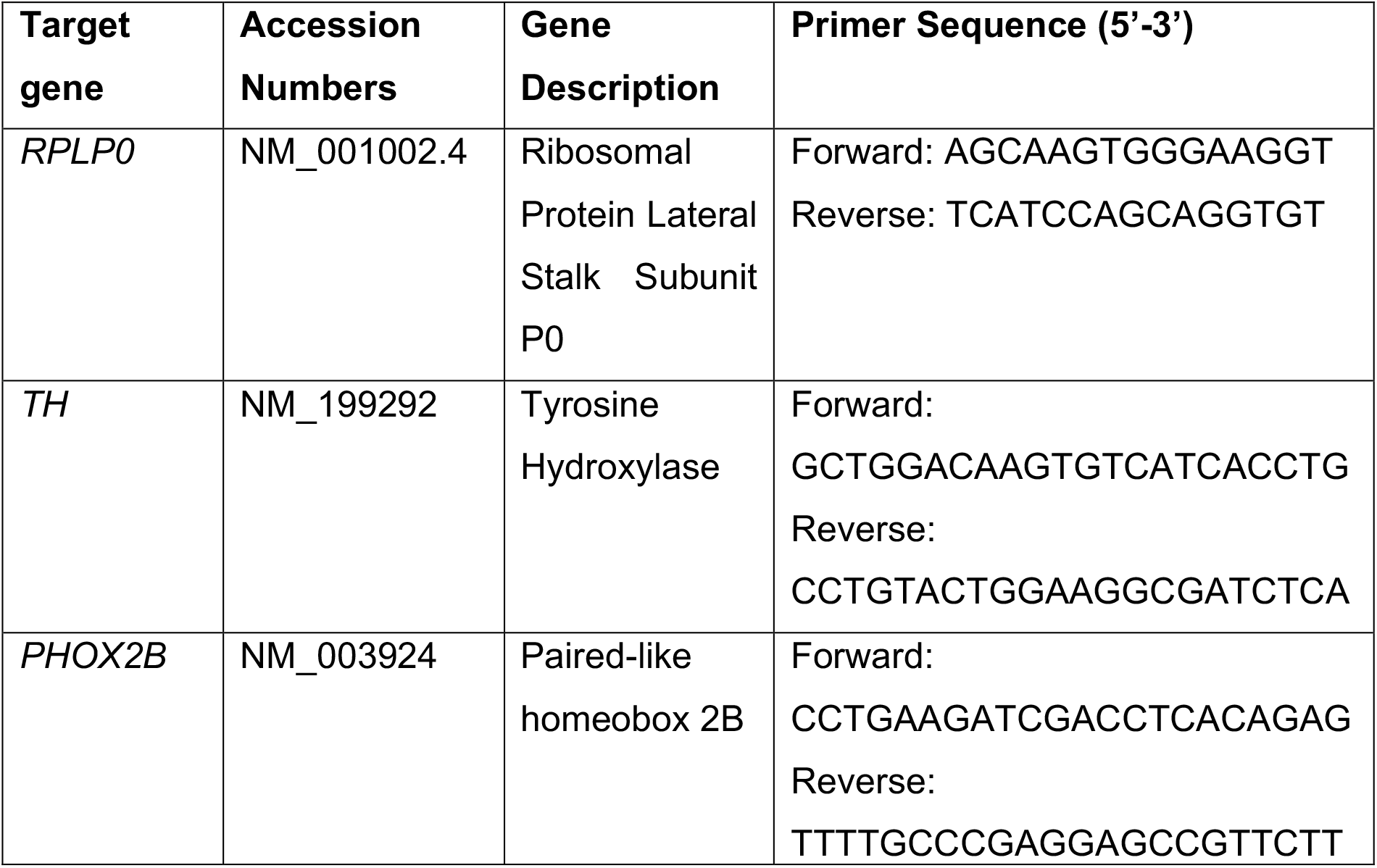
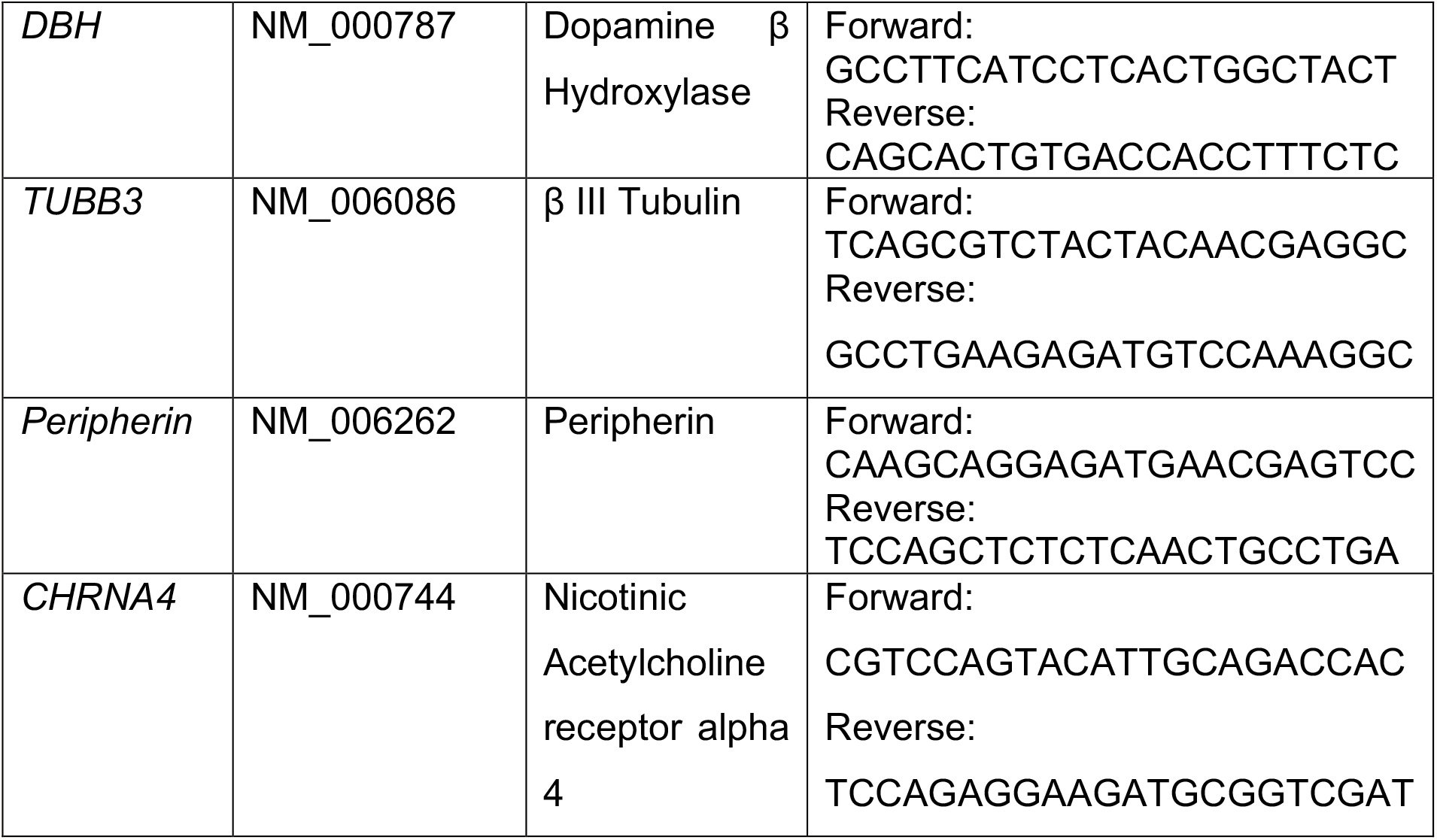
Summary of genes evaluated. Target gene, accession numbers, gene description and primer sequence.

### Whole cell electrophysiology

Patch clamp experiments were performed at room temperature using an Axopatch 200B amplifier (Axon Instruments, AZ, USA) as previously described^21^. Pipettes were made from borosilicate glass capillaries, and fire polished.

Spontaneous hiPSC-SNs APs were evaluated in gap free mode using the whole cell configuration. The patch pipettes (resistance 4-5 MOhms) were filled with a solution containing (in mmol/L): 120 K-Gluconate, 40 HEPES, 5 MgCl_2_, 2 Na-ATP and 0.3 Na-GTP; pH was adjusted at 7.2 with KOH. The bath solution (external current clamp) was composed of (in mmol/L): 119 NaCl, 2.5 KCl, 2.5 CaCl_2_, 1 Na_2_HPO_4_, 1.3 MgSO_4_, 26 NaHCO_3_, and 11 Glucose; the pH was adjusted at 7.4 with NaOH. During the recording, 1 µM Nicotine and 0.1 µM α-Bungarotoxin, an agonist and antagonist, respectively, of nicotinic acetylcholine receptor alpha 4 were applied to study the hiPSC-SNs APs responses.

Spontaneous hiPSC-CMs APs were evaluated in gap free mode using the whole cell configuration. The patch pipettes (resistance 2-3 MOhms) were filled with a solution containing (in mmol/L): 10 NaCl, 122 KCl, 1 MgCl_2_, 1 EGTA and 10 HEPES; pH was adjusted at 7.3 with KOH. The bath solution (external current clamp) was composed of (in mmol/L): 154 NaCl, 5.6 KCl, 2 CaCl_2_, 1 MgCl_2_, 8 glucose and 10 HEPES; the pH was adjusted at 7.3 with NaOH. During the APs recording, 1 µM nicotine or 0.1 µM α-bungarotoxin was applied in the neuronal compartment, and 5 µM propranolol, a β-receptor blocker was applied in the cardiomyocytes compartment, to observe the responses in the hiPSC-CMs APs and assess the functional neuro-cardiac connection.

### hiPSC-CMs contraction analysis

The hiPSC-CMs contractile function was performed using custom-made video analysis software, as previously published^22,23^. The MFD was placed on the stage of an inverted microscope equipped with a 20x objective. The spontaneous beating of hiPSC-CMs syncytium in the cardiomyocytes compartment were acquired at 60 frames per second, for hiPSC-CMs monocultured or co-cultured. Videos were then processed: TIFF images were extracted, and contrasted particle displacement was tracked frame by frame for each video. The displacement of each contrasted particle was then processed through the time duration, resulting in a curve of the displacement as a function of time. Areas with similar contractile behavior were clustered and contractile parameters quantified by a Matlab script. Similar to hiPSC-CMs, nicotine, α-bungarotoxin, and propranolol were applied to observe the beating rate response of hiPSC-CMs syncytia.

### Data analysis and statistics

Patch clamp results were analyzed using Clampfit (pCLAMP v10; Molecular devices, CA, USA), respectively. The contractile function was analyzed using custom-written Matlab programs. The code is available upon request. A normality test (Agostino and Pearson omnibus normality test) was always used to determine whether data followed a normal distribution. Data processing and statistical analyses were conducted with Prism 10. Mean with s.e.m. was calculated from experiments obtained from at least three independent differentiations. One- or two-way ANOVA test was used to assess significance (ns p > 0.05; *p < 0.05, **p < 0.01, ***p < 0.001).

## Results

### Characterization and validation of hiPSC-SNs identity

Immunofluorescence and RT-qPCR were performed to assess the sympathetic phenotype of the neurons. After 30 days of culture, hiPSC-SNs presented a “shiny bead” around the soma which is a morphological feature of sympathetic neuronal (**Figure 2A**). hiPSC-SNs were positive for DAPI (**Figure 2B**), tyrosine hydroxylase (TH), a catecholamine enzyme (**Figure 2C**), β-III tubulin (TUBB3, **Figure 2D**), a microtubule protein in neurons (merged of 2C and 2D is shown in **Figure 2E**), and for peripherin (**Figure 2F**), an intermediate filament protein found in neurons of the peripheral nervous system. Interestingly, *Paired-like homeobox 2B* (*PHOX2B*), a critical gene for development and maturation of ANS; *TH, dopamine β hydroxylase* (*DBH*), *TUBB3, peripherin* and *nicotinic acetylcholine receptor alpha 4* (*CHRNA4*) were also widely expressed in hiPSC-SNs compared to hiPSC-CMs, where no expression was evident by RT-qPCR (**Figure 2G-L**) confirming the sympathetic phenotype of the neurons.

**Figure 2:**
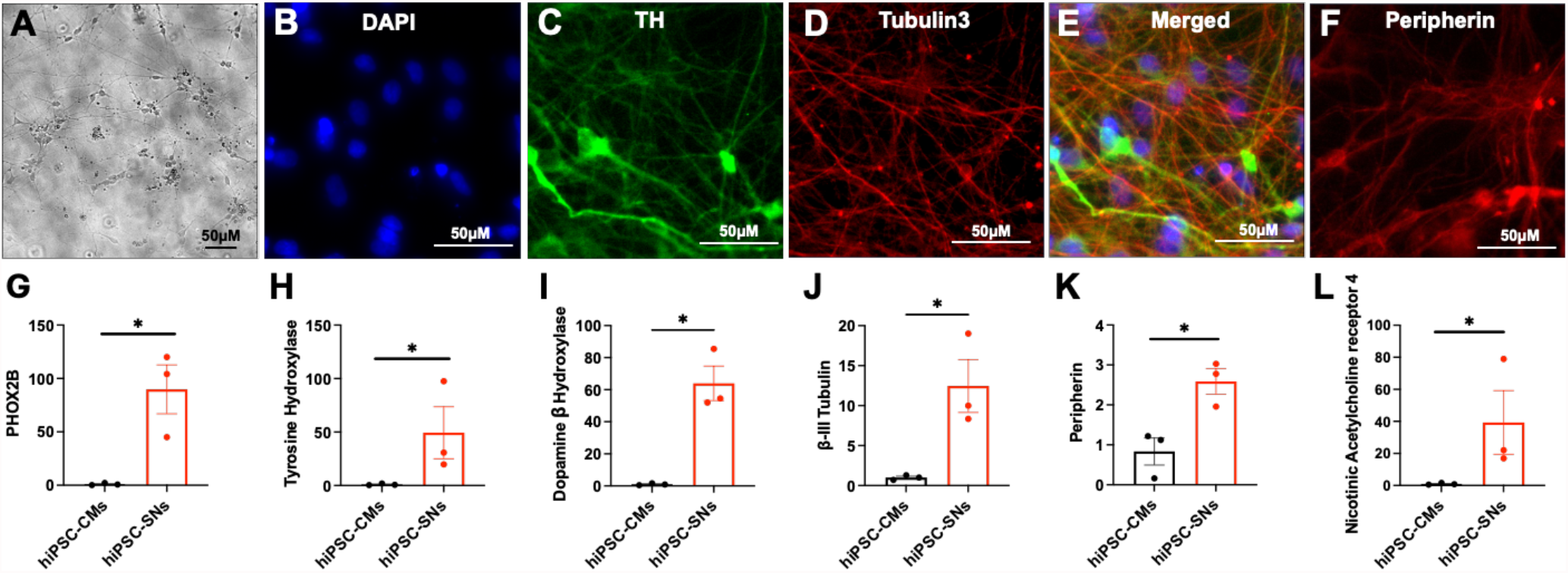
Characterization of sympathetic neurons-hiPSCs phenotype. **(A)** hiPSC-SNs observed at day 40 in transmitted light at x10 magnification, showing neuronal “shiny beads” morphology around the soma. **(B-F)** Immunohistochemical staining of day 40 hiPSC-SNs, positive for DAPI, the sympathetic neuronal enzyme tyrosine hydroxylase (TH), for microtubule protein in neurons β-III tubulin (TUBB3), and for the intermediate filament protein found in peripheric neurons (peripherin); observed at magnification X40. Fold change expression compared to hiPSC-CMs for **(G)** *Phox2B*, **(H)** *TH*, **(I)** *Dopamine β hydroxylase*, **(J)** *TUBB3*, **(K)** *Peripherin* and **(L)** *nicotinic acetylcholine receptor 4*. N=3 different neuronal differentiation; t-test Mann-Whitney. means ± s.e.m; *p < 0.05. hiPSC-CMs: hiPSC-cardiomyocytes; hiPSC-sympathetic neurons

We then performed patch-clamp experiments (**Figure 3A**) to assess the abilities of hiPSC-SNs to fire spontaneous or evoked APs. hiPSC-SNs exhibited spontaneous electrical activity manifesting as either a single or multiple APs (**Figure 3B**). Furthermore, in response to a long pulse (500ms), 90% of hiPSC-SNs displayed a tonic and accommodating firing pattern (**Figure 3C**). Collectively, these results characterize the functional electrophysiological profile of hiPSC-SNs. We then incubated the hiPSC-SNs with 0.1µM α-bungarotoxin and observed the responses of neuronal spontaneous APs firing. α-bungarotoxin, an antagonist of nicotinic acetylcholine receptor, alone could not slow the APs firing pattern of hiPSC-SNs (60 ± 3 to 59 ± 5 APs/min) (**Figure 3D**). We then applied an agonist of nicotinic acetylcholine receptor, 1 µM nicotine, which significantly enhanced the firing pattern (60 ± 3 to 97 ± 4 APs/min) (**Figure 3E**). Subsequent application of α-bungarotoxin did not affect this parameter, confirming the sympathetic response to pharmacological compound (**Figure 3F**).

**Figure 3:**
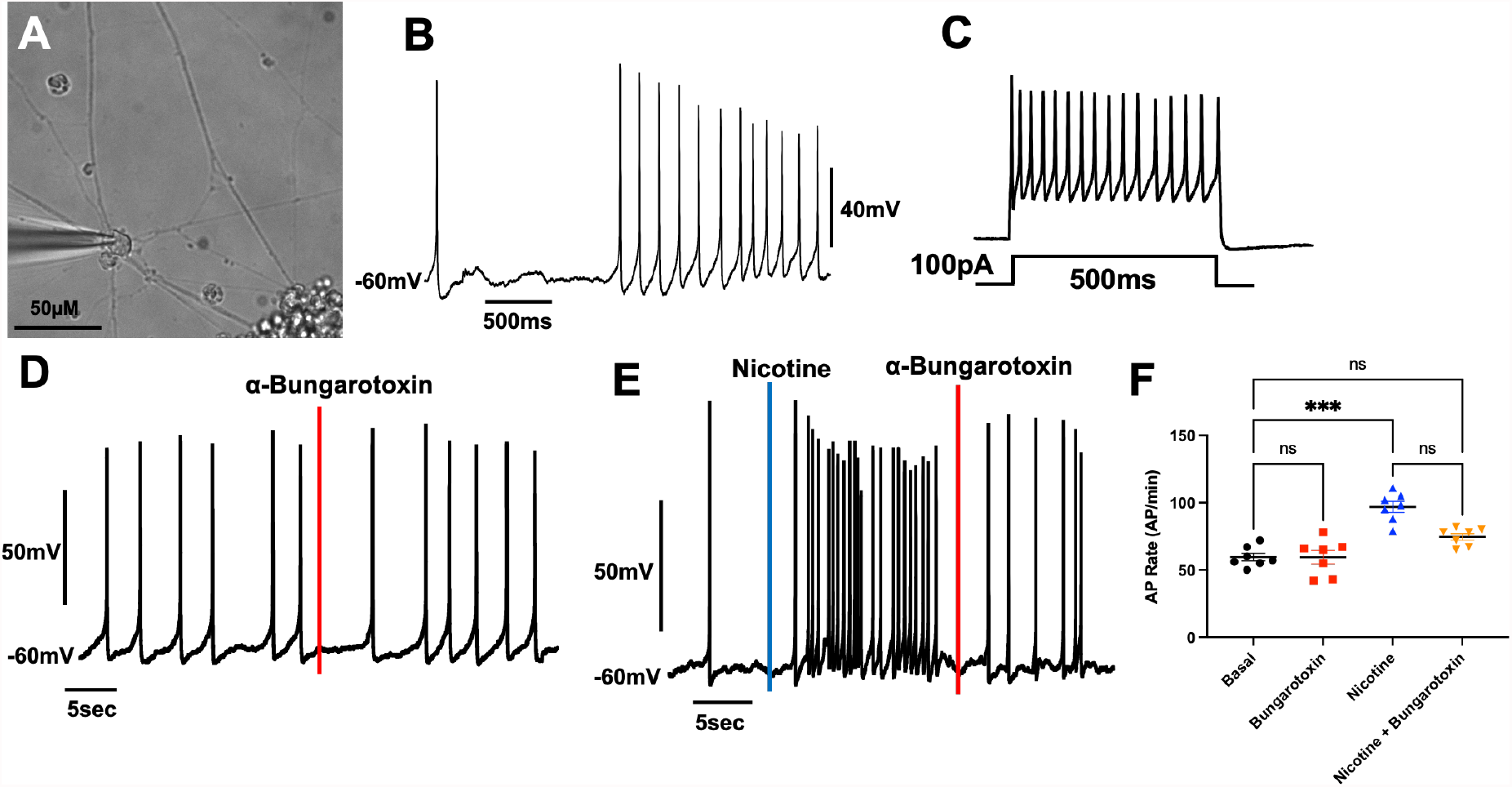
Electrophysiological validation of sympathetic neurons-hiPSCs. **(A and B)** Spontaneous action potential firing a single or multiple action potentials at day 40 for hiPSC-SNs in gap-free mode. **(C)** Tonic response could be evoked by a current injection duration of 500ms, showing action potentials accommodation in all hiPSC-SNs recorded. **(D)** Spontaneous action potentials were not affected by a sympathetic antagonist of nicotinic acetylcholine receptor, 0.1 µM α-bungarotoxin. **(E)** Spontaneous action potentials firing was increased by sympathetic agonist of nicotinic acetylcholine receptor, 1 µM nicotine, but less effects were observed after α-bungarotoxin application. **(F)** Action potential rate calculation illustrated by individual values scatter plot, validating the significant effect of nicotine in action potential rate. (n=7 for each condition, N=2 different neuronal differentiation). Kruskal-Wallis multiple comparison test; means ± s.e.m; ns: non-significant; ***p < 0.001.

### Co-culture of hiPSC-SNs and hiPSC-CMs in Microfluidic Device

After successfully differentiating hiPSCs into hiPSC-SNs and established positivity for the catecholaminergic marker and response to nicotine stimulation, we seeded hiPSC-SNs in the neuronal chamber of the MFD. After 7-10 days of culture, the neurites of the hiPSC-SNs successfully elongated through the microchannels and invaded the cardiomyocytes compartment (**Figure 4A and B**). When the neurites invaded the cardiomyocytes chamber, we seeded the hiPSC-CMs. After 10 days of co-culture, hiPSC-CMs syncytia (**Figure 4C**, red arrows) was connected to the neurites (**Figure 4C**, white arrow) and beating spontaneously. We also assess the physical connection between hiPSC-SNs and hiPSC-CMs by immunofluorescence. Our results confirmed that hiPSC-SNs seeded in the neuronal chamber (**Figure 4D**) marked with *TUBB3* prolonged neurites through the microchannels, and reached the hiPSC-CMs (**Figure 4E**) marked with α-actinin. The functional connection was then assessed by patch-clamp and video-recording.

**Figure 4:**
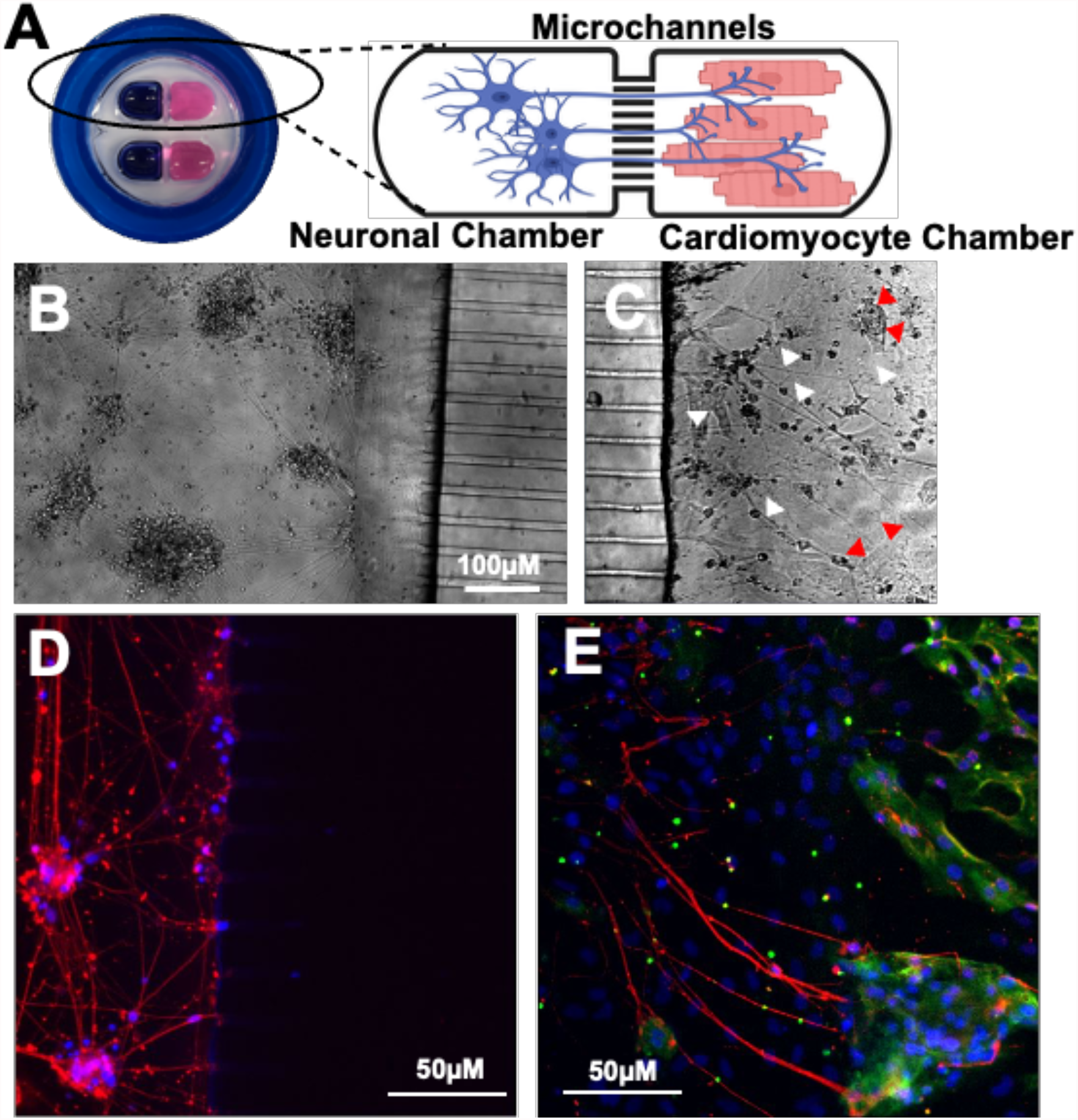
Microphysiological system to establish compartmentalized sympathetic neurons-cardiac junction on microfluidic device. **(A)** Representation of the microfluidic device, composed of two independent neuro-cardiac co-culture, with one neuronal chamber connected to cardiomyocyte chamber by microchannels. Neuronal observation in transmitted light in neuronal compartment **(B)** with neurites going to cardiomyocytes chamber **(C)**; white arrows showing the axons connected to hiPSC-CMs syncytium illustrated with red arrows. **(D and E)** Corresponding immunohistochemical staining of hiPSC-SNs (red=peripherin) connected to hiPSC-CMs (green=actinin); blue corresponding to DAPI; Magnification x20.

### Establishment of functional connection between hiPSC-SNs and hiPSC-CMs

The spontaneous activity was assessed by patch-clamp techniques in hiPSC-CMs in monoculture and co-cultured with hiPSC-SNs. The wide open-chamber of the MFD offer the opportunity to perform these experiments (**Figure 5A and B**). In monoculture, the application of 1 µM nicotine did not affect the spontaneous AP rate (20 ± 2 to 18 ± 2 AP/min), while the application of 5 µM propranolol reduced significantly the AP rate (20 ± 2 to 7 ± 1 AP/min) (**Figure 5C-F**). Interestingly, hiPSC-CMs exhibited a higher spontaneous APs rate when co-cultured in MFD compared to monoculture (20 ± 2 to 36 ± 3 AP/min) (**Figure 5G**). This time, the application of 1 µM of nicotine in the neuronal chamber significantly increase the AP rate (36 ± 3 to 74 ± 6 AP/min) (**Figure 5H**). Following the administration of nicotine, the addition of 5 µM propranolol to the cardiomyocyte chamber reduced the AP rate (**Figure 5I**). However, the values remained significantly higher than those observed in co-culture alone (36 ± 3 to 47 ± 2 AP/min) (**Figure 5F**).

**Figure 5:**
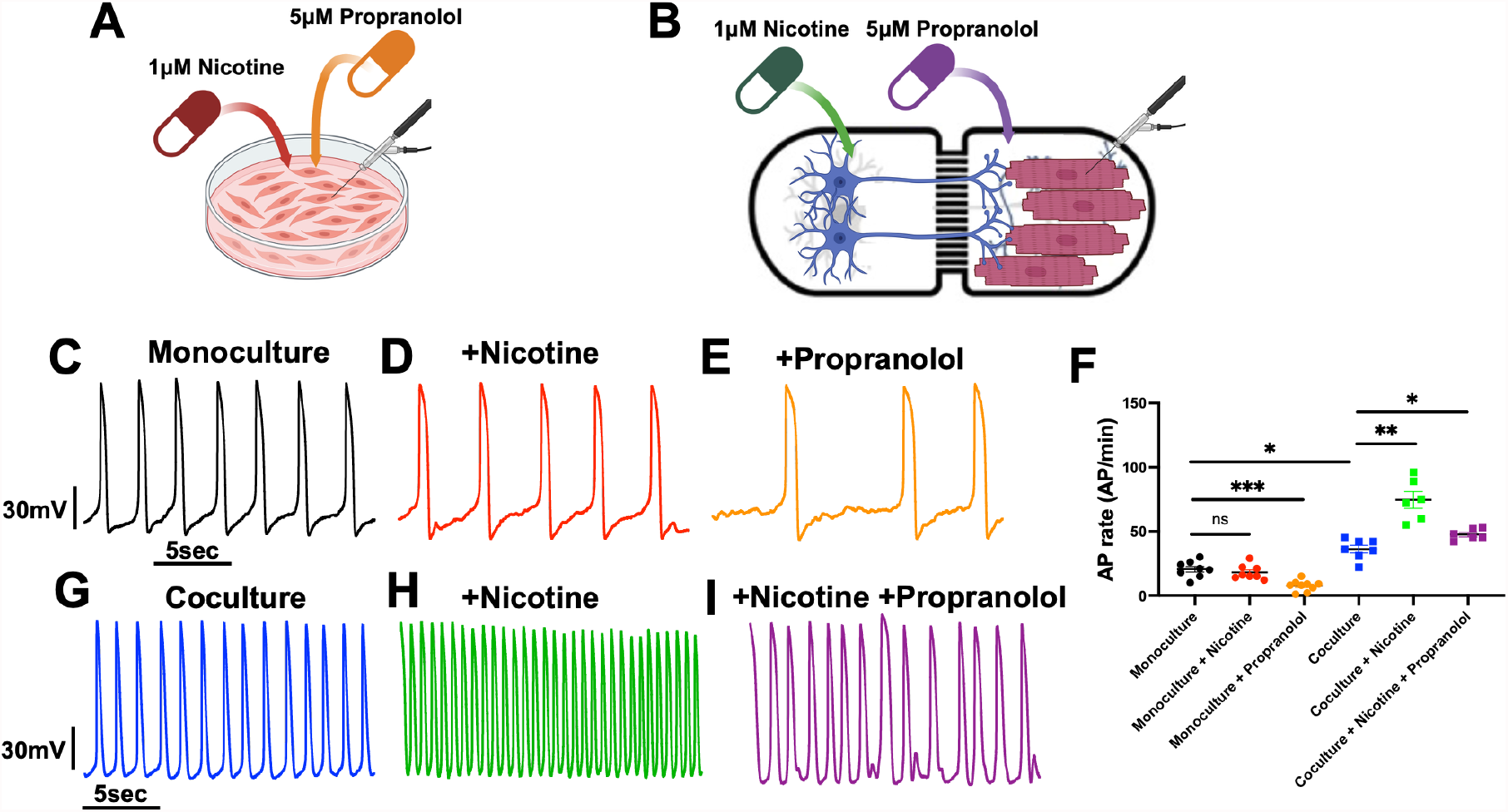
Sympathetic neurons-hiPSCs can control the spontaneous action potential rate of cardiomyocyte-hiPSCs in microfluidic device. **(A and B)** Schematic representation of patch-clamp experiments and pharmacological treatment in monoculture and co-culture. **(C-E)** Representative spontaneous action potentials firing in hiPSC-CM in monoculture (n=8), after 1µM nicotine application (n=8), and after 5µM propranolol application (n=10). **(F)** hiPSC-CMs action potential rate calculation illustrated by individual values scatter plot in monoculture and co-culture with hiPSC-SNs after pharmacological treatment. **(G-I)** Representative spontaneous action potentials firing in hiPSC-CM in co-culture with hiPSC-SNs (n=7), after 1µM nicotine application (n=6), and after 5µM propranolol application (n=6). Experiments were performed in N=2 different differentiation). Kruskal-Wallis multiple comparison test; means ± s.e.m; ns: non-significant; *p < 0.05, **p < 0.01, ***p < 0.001.

Next, we have recorded the motion properties of the syncytium hiPSC-CMs in monoculture *vs*. co-culture in MFD by video-recording. We used pharmacological compound to activate the hiPSC-SNs with nicotine or inhibited the β-adrenergic receptor in hiPSC-CMs with propranolol (**Figure 6A and B**). We evaluated the spontaneous beating rate of hiPSC-CMs syncytium in monoculture, where nicotine failed to induce changes in beating rate (33 ± 3 vs 28 ± 3 bpm), but propranolol significantly reduced the beating rate (33 ± 3 to 11 ± 2 bpm) (**Figure 6C-F**). Interestingly, in co-culture with hiPSC-SNs, the beating rate of hiPSC-CMs were significantly increased compared to hiPSC-CMs in monoculture (33 ± 3 to 58 ± 4 bpm) (**Figure 6F and G**). Application of nicotine significantly increased the beating rate, while propranolol normalized this increased, compared to co-culture in basal condition (58 ± 4 to 89 ± 3 to 53 ± 7 bpm) (**Figure 6F-I**). These experiments demonstrated a functional coupling between hiPSC-SNs and hiPSC-CMs in MFD.

**Figure 6:**
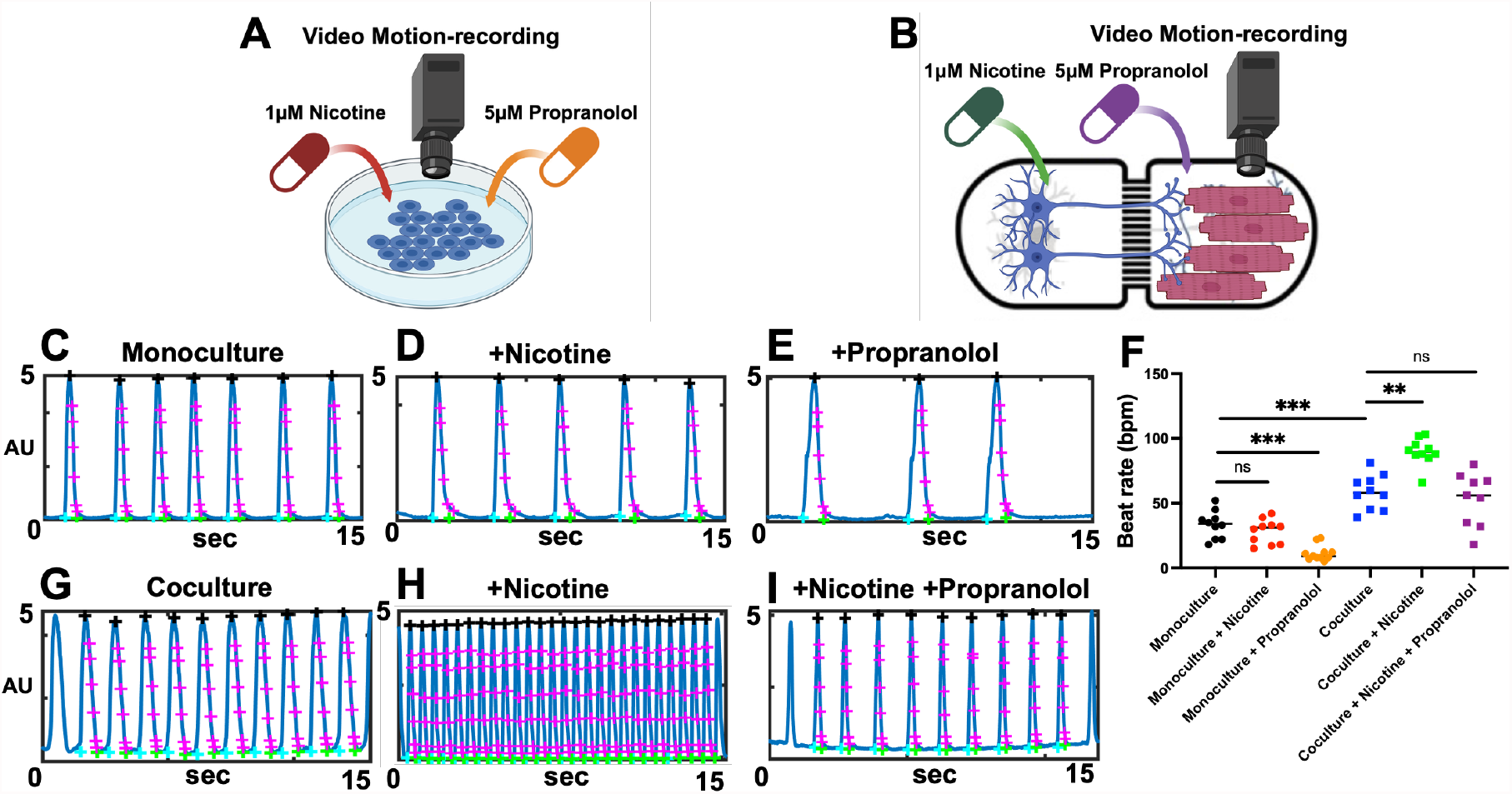
Sympathetic neurons-hiPSCs can control the beating rate of cardiomyocyte-hiPSCs in microfluidic device. **(A and B)** Schematic representation of video motion-recording experiments and pharmacological treatment in monoculture and co-culture. **(C-E)** Representative spontaneous beating rate of hiPSC-CMs in monoculture (n=10), after 1µM nicotine application (n=10), and after 5µM propranolol application (n=11). **(F)** hiPSC-CMs beating rate calculation illustrated by individual values scatter plot in monoculture and co-culture with hiPSC-SNs after pharmacological treatment. **(G-I)** Representative spontaneous beating rate of hiPSC-CM in co-culture with hiPSC-SNs (n=10), after 1µM nicotine application (n=10), and after 5µM propranolol application (n=9). Experiments were performed in N=2 different differentiation). Kruskal-Wallis multiple comparison test; means ± s.e.m; ns: non-significant; **p < 0.01, ***p < 0.001.

## Discussion

In this study, we created a MPS to investigate the functional impact of sympathetic innervation on cardiomyocytes from hiPSCs. We were able to differentiate hiPSC into hiPSC-SNs that exhibit catecholaminergic markers and express pro-catecholamine genes. In addition, these newly developed hiPSC-SNs exhibited a functional electrophysiology and a positive response to sympathetic agonist, nicotine. The co-culture between hiPSC-SNs and hiPSC-CMs in MFD demonstrated a physical and functional connection. Effectively, the spontaneous APs rate and beating rate of hiPSC-CMs were increased in basal condition and by the sympathetic agonist, nicotine, while the application of β-blocker, propranolol, decreased these parameters. The open-chamber architecture of the MFD used in this present paper, provide a unique compartmentalized platform for investigating how sympathetic innervation influences the electrophysiological and contractile properties of cardiomyocytes.

Our study is based on the use of hiPSCs differentiated into sympathetic neurons to create a MPS with hiPSC-CMs. We used previous protocols based on small molecule to differentiation the hiPSCs into neuromesodermal progenitor, trunk neural crest cell, and then neuronal precursors^18,24,25^. These previous studies, in addition to our data, demonstrated that the newly hiPSC-SNs formed by these neuronal precursor harbor catecholaminergic and adrenergic markers such as Phox2B, tyrosine hydroxylase, peripherin, and dopamine β hydroxylase. Therefore these markers are considered to be typical markers of sympathetic neurons^26^. In fact, in this neuron, the tyrosine is converted into L-DOPA by the tyrosine hydroxylase in the cytoplasm, then converted into dopamine by the DOPA decarboxylase, and finally converted into noradrenaline by the dopamine β hydroxylase^27–29^.

Electrical activity of hiPSC-SNs was studied by patch-clamp techniques. These cells showed spontaneous electrical activity characterized by single or multiple APs. The neurons responded to a prolonged current injection with accommodating pattern. We also assessed the respond of these neurons pharmacologically. The application of an antagonist of nicotinic acetylcholine receptor failed to modified the firing pattern of neuronal APs, while the application of agonist of these receptors was able to induce an increase of APs rate^30,31^. The difference between the nicotine and α-bungarotoxin can be explained by the difference of nicotinic acetylcholine receptor subtype present in hiPSC-SNs. α-bungarotoxin has only effect on *CHRNA* subtype 7, while nicotine has effect on the other subtypes express in sympathetic neurons, such as subtype 4^32,33^. The diversity of *CHRNA* subtype and function in the ANS could explain the difference observed, where the nicotine will induce an increase of firing pattern, which contribute to release noradrenaline, while α-bungarotoxin have less effect on the neuronal transmission^34^.

Several studies have described the co-culture of sympathetic neurons and cardiomyocytes, but present limitations by mixing species^4,35^ or mixing the co-culture media^18,19,36^.

In this study, we created a MPS by seeding hiPSC-SNs and hiPSC-CMs in each separated chamber of the MFD. After 10 days of co-culture, the neurite projections of hiPSC-SNs were able to connect physically to the hiPSC-CMs. We then validated the functional connection and formation of synapses by electrophysiological experiments and the application of pharmacological compounds. Interestingly, the co-culture, at the basal condition, increased the spontaneous AP and beating rate of hiPSC-CMs. The release of noradrenaline by the hiPSC-CMs activate the β-adrenergic receptors, which stimulate the adenylyl cyclase to generate the second messenger 3’-5’-cyclic adenosine monophosphate (cAMP)^37^. This increase of cAMP was previously described to facilitate the opening of hyperpolarization-activated cyclic nucleotide-gated (HCN) and Cav1.3 channel, contributing to increase the chronotropic responses of cardiomyocytes^38^. The application of nicotine, which stimulate the hiPSC-SNs firing pattern, increase the release of noradrenaline, while this compound has no effect on hiPSC-CMs in monoculture. The application of a non-selective β-adrenergic blocker, propranolol, significantly decrease the response to the noradrenaline release by hiPSC-SNs^39^.

A limitation of our study is that while the hiPSC-SNs were fully characterized, we did not evaluate the purity of these neurons. Markers and pharmacology stimulation confirm the phenotype of these neurons; however, we did not measure the level of catecholamine release in the culture media and especially close to the connection with cardiomyocytes. Previous study demonstrated that the neurons could release acetylcholine and dopamine, raising the question of the purity of the hiPSC-SNs^35^. This limitation did not affect the ability of these neurons to modulate the cardiomyocytes beating rates. In fact, the synaptic connection between neurons and cardiomyocytes is very narrow intercellular gap, within this space, noradrenaline is selectively released via the polarized neuronal active zone^40^. This specialized architecture ensures high local noradrenaline concentrations from only a minimal number of molecules, thereby facilitating the efficient activation of cardiac β-adrenoceptors.

Finally, the use of MFD between neurons and cardiomyocytes from hiPSCs could improve the maturity of hiPSC-CMs, which are a major challenging of using this type of cardiomyocytes^41^. Effectively, the cardiac sympathetic neurons secrete neuropeptide Y (NPY), which were previously described to affect the calcium-dependent process^42^, and activate the transient outward potassium current (Ito)^43,44^. The β-adrenergic stimulation was also previously described to contribute to hiPSC-CMs functional maturation^4^.

In conclusion, we successfully generated a functional MPS integrating hiPSC-SNs and hiPSC-CMs to recapitulate the physiological interaction between the ANS and the heart. We characterized the electrophysiological properties and adrenergic phenotype of these hiPSC-SNs, confirming their maturation and functionality. Our microfluidic-based demonstrated both physical and functional connectivity between the two cell types, as evidenced by their response to sympathetic neurons agonist and β-blockers. The advantages of MFD are the ability to control the culture media in each chamber and apply specific pharmacological compound in one chamber and observe the effect in the other. The design of this MFD also offers a significant advantage over previous devices, which were constrained by limited readout capabilities such as the application of patch clamp technique.

Ultimately, this platform provides a robust tool for investigating the brain-heart axis, a critical factor in various cardiac arrhythmias, identify new functional markers, and provide a more integrated understanding of ANS role in cardiac arrhythmias. This approach could enhance pharmacological testing during drug development, and support precision medicine strategies.

## Data availability

The data supporting this article and other findings are available within the manuscript, figures, supplementary figures and from the corresponding author upon request.

## Acknowledgements

We would like to acknowledge our fundings from Biomedical Laboratory Research & Development Service of United States (U.S.) Department of Veterans Affairs, Office of Research and Development, Merit Review grant I01 BX002137 to MB; National Heart, Lung, and Blood Institute 1R01HL164415 to MB; US Department of Defense award number W81XWH-21-1-0424 to MB.

## Author contributions

J-B. R. and Y.S. designed the experiments, contributed to the discussion, and wrote the manuscript. J-B.R. and Y.S collected and analyzed the data. J-B.R., Y.S. and M.B. contributed to the discussion and reviewed/re-edited the manuscript.

